# High Throughput Functional Evaluation of KCNQ1 Decrypts Variants of Unknown Significance

**DOI:** 10.1101/223206

**Authors:** Carlos G. Vanoye, Reshma R. Desai, Katarina L. Fabre, Franck Potet, Jean-Marc DeKeyser, Daniela Macaya, Jens Meiler, Charles R. Sanders, Alfred L. George

## Abstract

**Background:** The explosive growth in known human gene variation presents enormous challenges to current approaches for variant classification that impact diagnosis and treatment of many genetic diseases. For disorders caused by mutations in cardiac ion channels, such as congenital long-QT syndrome (LQTS), *in vitro* electrophysiological evidence has high value in discriminating pathogenic from benign variants, but these data are often lacking because assays are cost-, time- and labor-intensive.

**Methods and Results:** We implemented a strategy for performing high throughput, functional evaluations of ion channel variants that repurposed an automated electrophysiology platform developed previously for drug discovery. We demonstrated success of this approach by evaluating 78 variants in *KCNQ1*, a major LQTS gene. We benchmarked our results with traditional electrophysiological approaches and observed a high level of concordance. Our results provided functional data useful for classifying ~70% of previously unstudied *KCNQ1* variants annotated with uninformative descriptions in the public database ClinVar. Further, we show that rare and ultra-rare *KCNQ1* variants in the general population exhibit functional properties ranging from normal to severe loss-of-function indicating that allele frequency is not a reliable predictor of channel function.

**Conclusions:** Our results illustrate an efficient and high throughput paradigm linking genotype to function for a human cardiac channelopathy that will enable data-driven classification of large numbers of variants and create new opportunities for precision medicine.

## INTRODUCTION

The widespread use of genetic and genomic testing in research and clinical medicine has led to explosive growth in the number of gene variants associated with human diseases and in reference populations. This has led to an emerging challenge to classify variants accurately with respect to potential pathogenicity. The interpretation of clinical genetic tests are often confounded by ‘variants of unknown significance’ (VUS), a term indicating that there are insufficient data to distinguish between disease-causing mutations and benign variants.^1^ Several computational tools have been developed to predict which coding sequence variants may be deleterious,^2^ but these methods have not been systematically validated and results from such analyses are deemed to have lower value than experimental evidence for classifying variants in the clinical setting.^3^ There has been a recent call for expanded efforts to functionally annotate variants,^4^ but not all genes are equally amenable to large scale functional studies.

Human genetic diseases caused by mutations in ion channels (channelopathy)^5^ represent unique opportunities to meet the challenge of variant annotation because well-established *in vitro* functional assay paradigms exist for these proteins. The standard approach for determining the functional properties of an ion channel variant is cellular electrophysiology using patchclamp recording of heterologously expressed recombinant channels. However the usual embodiment of this method is tedious, time- and labor-intensive, making it too low throughput for determining the functional consequences of more than a few variants at a time.

One channelopathy that illustrates the challenge of variant classification is the congenital long-QT syndrome (LQTS), an inherited predisposition to sudden cardiac death affecting children and young adults ^6–8^ with a population incidence of approximately 1 in 2500.^9^ As with many other inherited disorders, clinical genetic testing has become standard-of-care for LQTS.^10,11^ Nearly 50% of LQTS cases are associated with genetic variants in *KCNQ1*^12^ that encodes the voltage-gated potassium channel K_V_7.1, which complexes with an accessory subunit encoded by *KCNE1* to generate the slow delayed rectifier current (*I*_Ks_) essential for normal myocardial repolarization.^13,14^ In addition to LQTS, at least two other genetic arrhythmia syndromes, short QT syndrome^15,16^ and familial atrial fibrillation,^17,18^ as well as cases of sudden infant death syndrome^19–21^ are associated with *KCNQ1* mutations. There are currently more than 600 known disease-associated *KCNQ1* variants,^22^ but only a small fraction has been analyzed functionally. Further, there are many rare *KCNQ1* variants found in population genome sequence databases, some of which have minor allele frequencies lower than the estimated population prevalence of LQTS. Accurately discriminating pathogenic *KCNQ1* mutations from benign variants has actionable consequences for diagnosis, treatment, prognosis and family counseling. Functional annotation of *KCNQ1* variants would offer strong supporting evidence for variant classification.

To overcome the challenge of determining the function of hundreds of ion channel variants, we coupled two advanced technologies: high efficiency cell electroporation and automated planar patch clamp recording. In this study, we demonstrated the capabilities of this approach by investigating the functional consequences of 78 *KCNQ1* variants parsed into training (30 variants) and test (48 variants) sets. Our study offers a new paradigm for the functional evaluation of ion channel gene variants using a high throughput experimental approach and represents a milestone in the application of automated cellular electrophysiology.

## METHODS

### Cell culture

Chinese hamster ovary cells (CHO-K1, CRL 9618, American Type Culture Collection, Manassas VA, USA) were grown in F-12 nutrient mixture medium (GIBCO/Invitrogen, San Diego, CA, USA) supplemented with 10% fetal bovine serum (ATLANTA Biologicals, Norcross, GA, USA), penicillin (50 units·ml^−1^), streptomycin (50 μg·ml^−1^) at 37°C in 5% CO_2_. Unless otherwise stated, all tissue culture media was obtained from Life Technologies, Inc. (Grand Island, NY, USA).

### Plasmids and mutagenesis

Full-length cDNA encoding human KCNQ1 (GenBank accession AF000571) or KCNE1 (GenBank accession L28168) were engineered in the mammalian expression vectors pIRES2-EGFP (BD Biosciences-Clontech, Mountain View, CA, USA) and a modified pIRES2-vector in which we replaced the fluorescent protein coding sequence with that of DsRed-MST (pIRES2-DsRed-MST, KCNE1) as previously described.^23,24^

Variants in the KCNQ1 voltage sensor domain (VSD, residues 100-250) were introduced using QuikChange II XL (Agilent technologies, Santa Clara, CA, USA). Mutagenic primers (see Supplemental Table S4 for sequences) for each variant were designed using the online QuikChange primer design tool (http://www.genomics.agilent.com/primerDesignProgram.jsp). Variant KCNQ1 plasmid clones were screened in batches of 12. Briefly, 96 colonies (12 variants × 8 clones) were picked and inoculated into LB broth supplemented with kanamycin (25 μg/ml) in a 96-well, deep-well plate (Cat # AB-0661, Thermo Fisher Scientific, Waltham, MA, USA) and incubated shaking (335 rpm) at 37^o^C for 16-18 h. The plate was then centrifuged at 1800 × g for 30 mins in a swing-out rotor (Allegra 6KR, Beckman Coulter, Indianapolis, IN, USA). The supernatant was discarded and the cell pellet in each well was processed using commercial alkaline lysis reagents (Macherey-Nagel Inc., Bethlehem, PA, USA). Following the final step of plasmid isolation, the plate was centrifuged at 1800 × g for 30 minutes and 200 μl of supernatant was transferred into a 96-well filter plate (Cat. # MSDVN6510, EMD Millipore, Billerica, MA, USA) secured on top of a 96-well collection plate (0.65 mL deep well plate, Thermo-Fisher Scientific) pre-filled with 330 μl of 100% ethanol. The stacked filter/deep well plates were spun at 1800 × g for 30 minutes to precipitate DNA, which was then washed once with 70% ethanol, dried and redissolved in TE buffer (10 mM Tris, 1 mM EDTA, pH 8.0).

Complete coding regions of KCNQ1 plasmids were sequenced (Eurofins Genomics, Louisville, KY) and analyzed using a custom multiple sequence alignment tool (MuSIC - Multiple Sequence Iterative Comparator, source code available upon request). Plasmid DNA from clones with correct sequences was amplified using an endotoxin-free plasmid prep method (Nucleobond Xtra Maxi EF, Macherey-Nagel Inc.) and re-suspended in endotoxin-free water.

### Electroporation

Plasmids encoding wild-type (WT) KCNE1 and KCNQ1 (WT or variants) were transiently co-transfected into CHO-K1 cells by electroporation using the Maxcyte STX system (MaxCyte Inc., Gaithersburg, MD, USA). CHO-K1 cells grown to 70-80% confluency were harvested using 5% trypsin. A 500 μl aliquot of cell suspension was then used to determine cell number and viability on an automated cell counter (ViCell, Beckman Coulter). Remaining cells were collected by gentle centrifugation (160 × g, 4 minutes), followed by washing the cell pellet with 5 ml electroporation buffer (EBR100, MaxCyte Inc.) and re-suspension in electroporation buffer at a density of 10^8^ viable cells/ml.

For each electroporation, plasmids encoding WT KCNE1 (30 μg) and individual KCNQ1 variants (15 μg) were mixed and then added to 100 μl cell suspension (10^8^ cells/ml). The DNA-cell suspension mix was then transferred to an OC-100 processing assembly (MaxCyte Inc.) and electroporated using the CHO-PE preset protocol. Immediately after electroporation, 10 μl of DNase I (Sigma-Aldrich, St. Louis, MO, USA) was added to the DNA-cell suspension and the entire mixture was transferred to a 35 mm tissue culture dish and incubated for 30 min at 37°C in 5% CO_2_. Following incubation, cells were gently re-suspended in culture media, transferred to a T75 tissue culture flask and grown for 48 hours at 37°C in 5% CO_2_. Following incubation, cells were harvested, counted, transfection efficiency determined by flow cytometry (see below) and then frozen in 1 ml aliquots at 1.5×10^6^ viable cells/ml in liquid N_2_ until used in experiments.

### Flow Cytometry

Transfection efficiency was evaluated before freezing and prior to electrophysiological testing using a benchtop flow cytometer (CytoFLEX, Beckman Coulter). Forward scatter (FSC), side scatter (SSC), green fluorescence (FITC), and red fluorescence (PE) were recorded. FSC and SSC were used to gate single viable cells and to eliminate doublets, dead cells and debris. Ten thousand events were recorded for each sample. Non-electroporated CHO-K1 cells were assayed as a control for all parameters and used to set the gates for each experiment. A 488 nm laser was used to excite both FITC and PE, and the percentage of fluorescent cells was determined from the gated population. A compensation matrix was generated using CHO-K1 cells expressing only GFP or dsRed fluorescent markers and applied to the co-transfected cells to account for spectrum overlap. The percentage of co-transfected cells was determined from plots of FITC *vs* PE fluorescence intensity.

### Cell preparation for automated electrophysiology

Co-electroporated cells were thawed the day before experiments, plated and incubated for 10 hours at 37°C in 5% CO_2_. The cells were then transferred to 28°C in 5% CO_2_ and grown overnight. Prior to the experiment, cells were passaged using 5% trypsin in cell culture media. Cell aliquots (500 μl) were used to determine cell number and viability by automated cell counting while transfection efficiency was determined by flow cytometry. Cells were then diluted to 200,000 cells/ml with external solution (see below), and allowed to recover 40 minutes at 15°C while shaking on a rotating platform at 200 rpm.

### Automated patch clamp recording

Automated patch clamp recording was performed using the Syncropatch 768 PE platform (Nanion Technologies, Munich, Germany). Single-hole, 384-well recording chips with medium resistance (2-4 MΩ) were used in this study. Pulse generation and data collection were carried out with PatchController384 V.1.3.0 and DataController384 V1.2.1 software (Nanion Technologies). Whole-cell currents were filtered at 3 kHz and acquired at 10 kHz. The access resistance and apparent membrane capacitance were estimated using built-in protocols. Whole-cell currents were recorded at room temperature in the whole-cell configuration from −80 to +60 mV (in 10 mV steps) 1990 ms after the start of the voltage pulse from a holding potential of −80 mV. The external solution contained (in mM) the following: NaCl 140, KCl 4, CaCl_2_ 2, MgCl_2_ 1, HEPES 10, glucose 5, with the final pH adjusted to 7.4 with NaOH. The internal solution contained (in mM) the following: KF 60, KCl 50, NaCl 10, HEPES 10, EGTA 10, ATP-K_2_ 5, with the final pH adjusted to 7.2 with KOH. Whole-cell currents were not leak-subtracted. The contribution of background currents was determined by recording before and after addition of the I_Ks_ blocker HMR1556 (20 μM). Only HMR1556-sensitive currents were used for analysis. Stringent criteria were used to include individual cell recordings for final data analysis (seal resistance ≥ 0.5 gigaohm; series resistance ≤ 20 megaohm; capacitance ≥ 1 picofarad; voltage-clamp stability as indicated by a standard error for baseline current measured at the holding potential for all test pulses being <10% of the mean baseline current). Current-voltage relationships were derived for all cell recordings that met these criteria. Unless otherwise stated, all chemicals were obtained from (Sigma-Aldrich, St. Louis, MO, USA).

### Manual patch clamp recording

Whole-cell currents were recorded at room temperature (RT, 20−23°C) using Axopatch 200 and 200B amplifiers (Molecular Devices Corp., Sunnyvale, CA, USA) in the whole-cell configuration of the patch clamp technique.^25^ Pulse generation was performed with Clampex 10.0 (Molecular Devices Corp.). Whole-cell currents were filtered at 1 kHz and acquired at 5 kHz. The access resistance and apparent membrane capacitance were estimated using an established method.^26^ Whole-cell currents were not leak-subtracted. Whole-cell currents were measured from −80 to +60 mV (in 10 mV steps) 1990 ms after the start of the voltage pulse from a holding potential of −80 mV. The external solution contained (in mM) the following: NaCl 132, KCl 4.8, MgCl_2_ 1. 2, CaCl_2_ 1, glucose 5, HEPES 10, pH 7.4. The internal solution contained (in mM) the following: K+ aspartate 110, CaCl_2_ 1, HEPES 10, EGTA 11, MgCl_2_ 1, ATP-K_2_ 5, pH 7.3 adjusted with KOH. Pipette solution was diluted 5-10% to prevent activation of swelling-activated currents. Patch pipettes were pulled from thick-wall borosilicate glass (World Precision Instruments, Inc., Sarasota, FL, USA) with a multistage P-97 Flaming-Brown micropipette puller (Sutter Instruments Co., San Rafael, CA, USA) and heat-polished with a Micro Forge MF 830 (Narashige, Japan). After heat polishing, the resistance of the patch pipettes was 1.5-3 MΩ in the standard extracellular solution. The access resistance varied from 3 to 9 MΩ and experiments were excluded from analysis if the voltage errors originating from the series resistance were greater than 5 mV. As a reference electrode, a 2% agar-bridge with composition similar to the control bath solution was utilized. Junction potentials were zeroed with the filled pipette in the bath solution. Unless otherwise stated, all chemicals were obtained from (Sigma-Aldrich, St. Louis, MO, USA).

### Data analysis

Data were analyzed and plotted using a combination of DataController384 V1.2.1 (Nanion Technologies, Clampfit V10.4 (Molecular Devices Corp.), Excel (Microsoft Office 2013, Microsoft), SigmaPlot 2000 (Systat Software, Inc., San Jose, CA USA) and OriginPro 2016 (OriginLab, Northampton, MA). Additional custom semi-automated data handling routines were used for rapid analysis of current density, voltage-dependence of activation and gating kinetics. Whole-cell currents were normalized for membrane capacitance and results expressed as mean ± SEM. The voltage-dependence of activation was determined only for cells with mean current density values greater than the background current amplitude. The number of cells used for each experimental condition is given in the figure legends and tables.

## RESULTS

### Coupling high efficiency cell electroporation to automated electrophysiology

Unlike manual patch clamp recording, which affords an opportunity to select cells by microscopy, automated electrophysiology is performed without direct visualization. This precludes the typical use of fluorescent or other co-transfection markers to identify cells at the time of experiments. Generating cell lines stably expressing a protein of interest is the usual solution to this problem, but substantial time and cost barriers are incurred when analyzing hundreds of variants. To overcome this obstacle, we implemented a high efficiency cell electroporation method for transiently co-expressing plasmids encoding KCNQ1 and its accessory subunit, KCNE1, suitable for automated electrophysiology (see Methods). We expressed channel subunits from plasmid vectors encoding either green (EGFP coupled to KCNQ1) or red (DsRed coupled to KCNE1) fluorescent proteins to enable quantification of transfection efficiency by flow cytometry.

Optimized co-transfection efficiency and cell survival were obtained by electroporating 30 μg KCNE1 with 15 μg KCNQ1 plasmids into Chinese hamster ovary (CHO-K1) cells. Under these conditions, we obtained 88.2 ± 5.4% (mean ± S.D.) transfection efficiency for WT KCNQ1 (inferred by the proportion of green fluorescent cells) and 81.9 ± 7.3% transfection efficiency for KCNE1 (red fluorescence) resulting in a co-transfection efficiency of 80.0 ± 8.2% with an average cell viability of 94.2 ± 6.8% (n = 32 transfections; Fig. 1). Non-transfected CHO-K1 cells lacked inherent green or red fluorescence (Supplementary Fig. S1), thus eliminating concern for false positives or over-estimation of transfection efficiencies by flow cytometry. Serial analysis demonstrated a high degree of reproducibility of these results. Cells were cultured for 48 h after electroporation then cryopreserved. Sufficient numbers of cells survived each electroporation to enable at least two automated electrophysiology experiments.

**Fig. 1.**
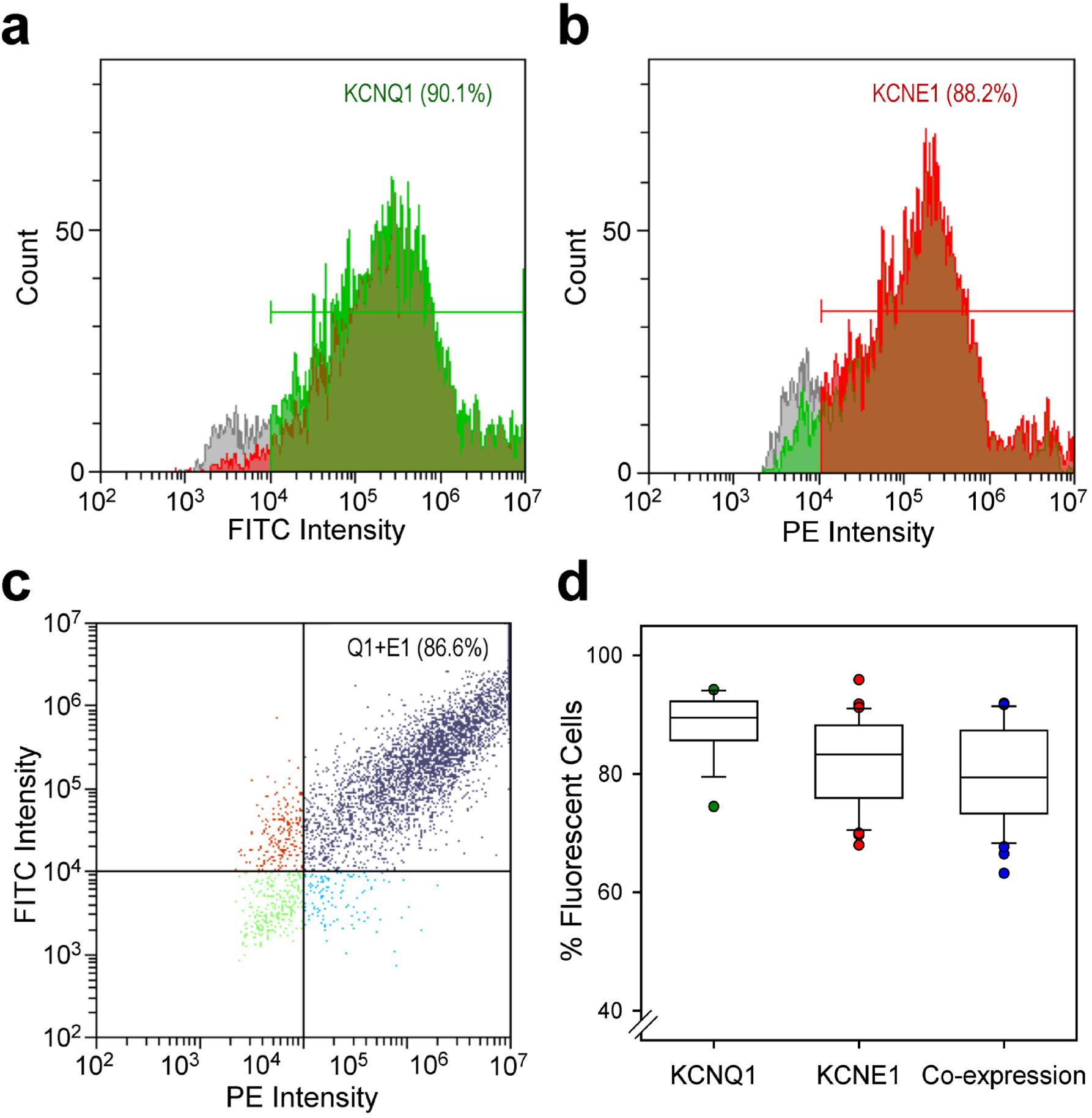
Efficiency of KCNQ1 and KCNE1 electroporation. Representative plots of fluorescence intensity *vs* cell counts measured by flow cytometry for CHO cells co-transfected with KCNQ1 (**a**, green fluorescence, FITC) and KCNE1 (**b**, red fluorescence, PE). The percentage of fluorescent cells is indicated within the panels. **c**) Plot of FITC *vs* PE intensity illustrating efficiency of KCNQ1 (Q1) and KCNE1 (E1) co-transfection (purple dots, upper right quadrant). **d**) Box plots illustrating the range of transfection efficiencies for KCNQ1, KCNE1 and co-expression for 32 separate electroporation experiments. Each box height represents the 25^th^ to 75^th^ percentile and the error bars represent the 5^th^ to 95^th^ percentile. Individual data points falling outside the 5-95^th^ percentile range are plotted. Solid and dotted horizontal lines within each box indicate the median and mean values, respectfully.

We recorded robust whole-cell current with properties consistent with *I*_Ks_ from cells electroporated with KCNQ1 and KCNE1 plasmids using an automated, dual 384-well planar patch clamp system (see Methods; Fig. 2a). To eliminate background currents, we applied the selective I_Ks_ blocker HMR1556^27,28^ at the end of each experiment and performed offline subtraction of HMR1556-insensitive current. In a typical experiment, cells were captured in ~96% of plate wells (blue and green shaded wells, Fig. 2a) and ~300 (80% of 384-well plate) cells had seal resistance ≥0.5 GΩ (green shaded wells, Fig. 2a). The averaged peak current density measured by automated patch clamp was lower than that measured by manual patch clamp (Fig. 2b). However, the averaged whole-cell currents normalized to the maximum peak current amplitude exhibited nearly identical gating kinetics measured by the two approaches (Fig. 2c) and had indistinguishable apparent half-maximal activation voltages (V_½_) (manual patch clamp: V_½_ = 23.0 ± 1.0 mV, n = 19; automated patch clamp: V_½_ = 24.5 ± 0.6 mV, n = 250; p = 0.5; Fig. 2d). The smaller currents recorded by automated patch clamp are partly explained by the unbiased selection of cells for recording in contrast to manual patch clamp in which the operator selects cells based on visually detectable fluorescence with a bias toward cells exhibiting bright fluorescence and, by inference, high channel expression. These results demonstrated that WT I_Ks_ recorded from electroporated cells by automated patch clamp is comparable to measurements made using manual patch clamp.

**Fig. 2.**
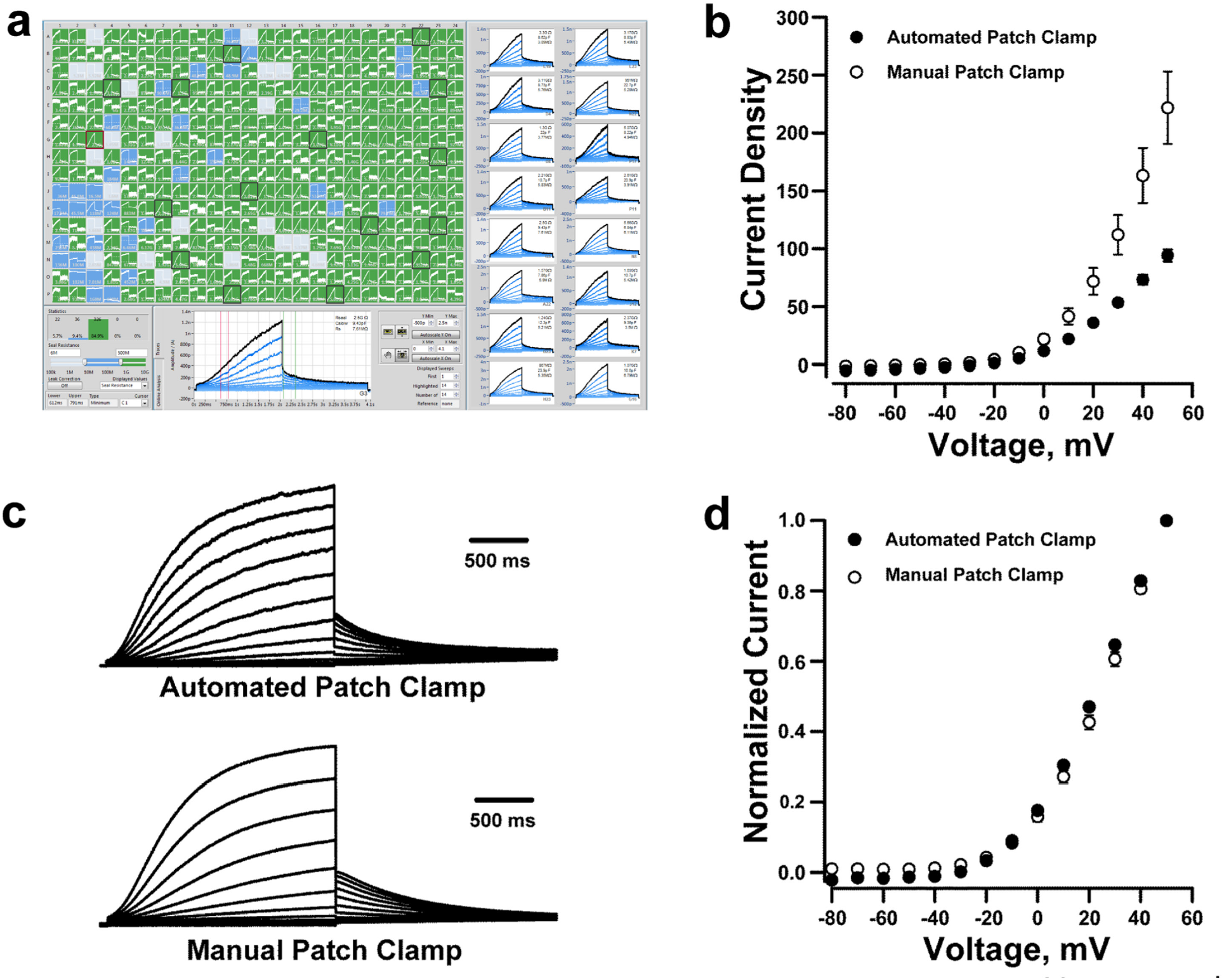
Automated patch clamp recording of wild type KCNQ1. **a**) Screen-shot of automated whole-cell current planar patch clamp recordings from CHO-K1 cells electroporated with plasmids encoding wild type (WT) KCNQ1 and KCNE1. The panels on the right highlight 16 out of the 384 wells that show whole-cell currents with I_Ks_-like kinetics. **b**) Current density-voltage relationships obtained from CHO-K1 cells transfected with KCNQ1 and KCNE1 recorded by automated (●, n = 250) or manual (◦, n = 19) patch clamp. Whole-cell currents are normalized by membrane capacitance (units are pA/pF). **c**) Average whole-cell currents recorded from cells electroporated with KCNQ1 and KCNE1 recorded using automated *(top)* or manual *(bottom)* patch clamp and normalized to the maximum peak current at +60 mV. **d**) Voltage-dependence of activation curves obtained from whole-cell currents recorded using automated (●) or manual (◦) patch clamp.

### Training and validation

We further validated and optimized throughput of the method using a training set of 30 disease-associated and non-disease associated KCNQ1 variants. The disease-associated variants included 17 variants (15 associated with LQTS; Supplemental Table S1) for which there were available functional data obtained using conventional (manual) electrophysiological approaches either from published reports or our laboratory. The non-disease associated set was comprised of 13 variants designed to match evolutionary substitutions found in orthologous KCNQ1 proteins from 12 different species (Supplemental Fig. S2), which we predicted would largely preserve channel function in contrast to many of the disease-associated variants that cause loss-of-function. Most of the variants chosen for this analysis affected residues located within the KCNQ1 voltage-sensing domain (VSD, residues 100-248). Variants were engineered into the KCNQ1 plasmid by site-directed mutagenesis and then electroporated into CHO-K1 cells prior to automated patch clamp recording experiments (see Methods).

A typical experiment consisted of a 384-well planar patch plate loaded with five distinct KCNQ1 variants, each seeded into 64 wells along with a separate cluster of wells with cells expressing either WT-KCNQ1 as a positive control or non-transfected cells as a negative control (average transfection efficiencies were 80.6 ± 6.8%, 74.2 ± 8.0% and 68.9 ± 7.6% for KCNQ1, KCNE1 and co-transfections, respectively; average viability was 88.8 ± 6.7%). Data from eight 384-well plates were sufficient to generate data for all 30 variants with dense replication (average number of successful replicates: n = 56 for variants, n = 301 for WT cells, and n = 220 for nontransfected cells). In these experiments, 94.4 ± 0.7% of wells exhibited cell capture with 80.4 ± 2.7% achieving a seal resistance of ≥ 0.5 GΩ yielding 1,976 cells that met stringent criteria and were used for analysis (see Methods).

Figure 3A illustrates averaged HMR1556-sensitive whole-cell current density recorded at +60 mV from CHO-K1 cells expressing the disease-associated (red) or evolutionary substitution (magenta) variants in the training set expressed as percent of WT, whereas Supplemental Fig. S3 illustrates averaged whole-cell current traces for all variants. Among the disease-associated KCNQ1 variants in the training set, most (12/17) generated current densities with small amplitudes (≤ 25% of WT) consistent with severe loss-of-function, which is the established molecular mechanism underlying LQTS. By contrast, 10 of the 13 evolutionary variants exhibited normal or near normal (defined as 70% to 130% of WT *I*_Ks_) current amplitudes, with 1 variant yielding current density >180% of WT and two others showing moderate (~50% of WT) loss- or gain-of-function. The voltage-dependence of activation was determined for all training set variants having a current amplitude >10% of WT and expressed as the divergence in half-maximal voltage (ΔV_½_) from WT (Fig. 3b). Comparisons of our findings obtained with automated patch clamp recording with published and unpublished data generated by manual patch clamp recording revealed a high degree of concordance (Fig. 4). A notable difference in activation ΔV_½_ between manual and automated patch clamp was observed for S209P likely because channels do not fully deactivate at the holding potential of −80 mV (Fig. 5) making the calculation of V_½_ unreliable by either recording method. Impaired deactivation of R231C also precluded reliable measurement of activation voltage-dependence. We did not determine a V_½_ value for L236R because the recorded currents were very small (~1% of WT).

**Fig. 3.**
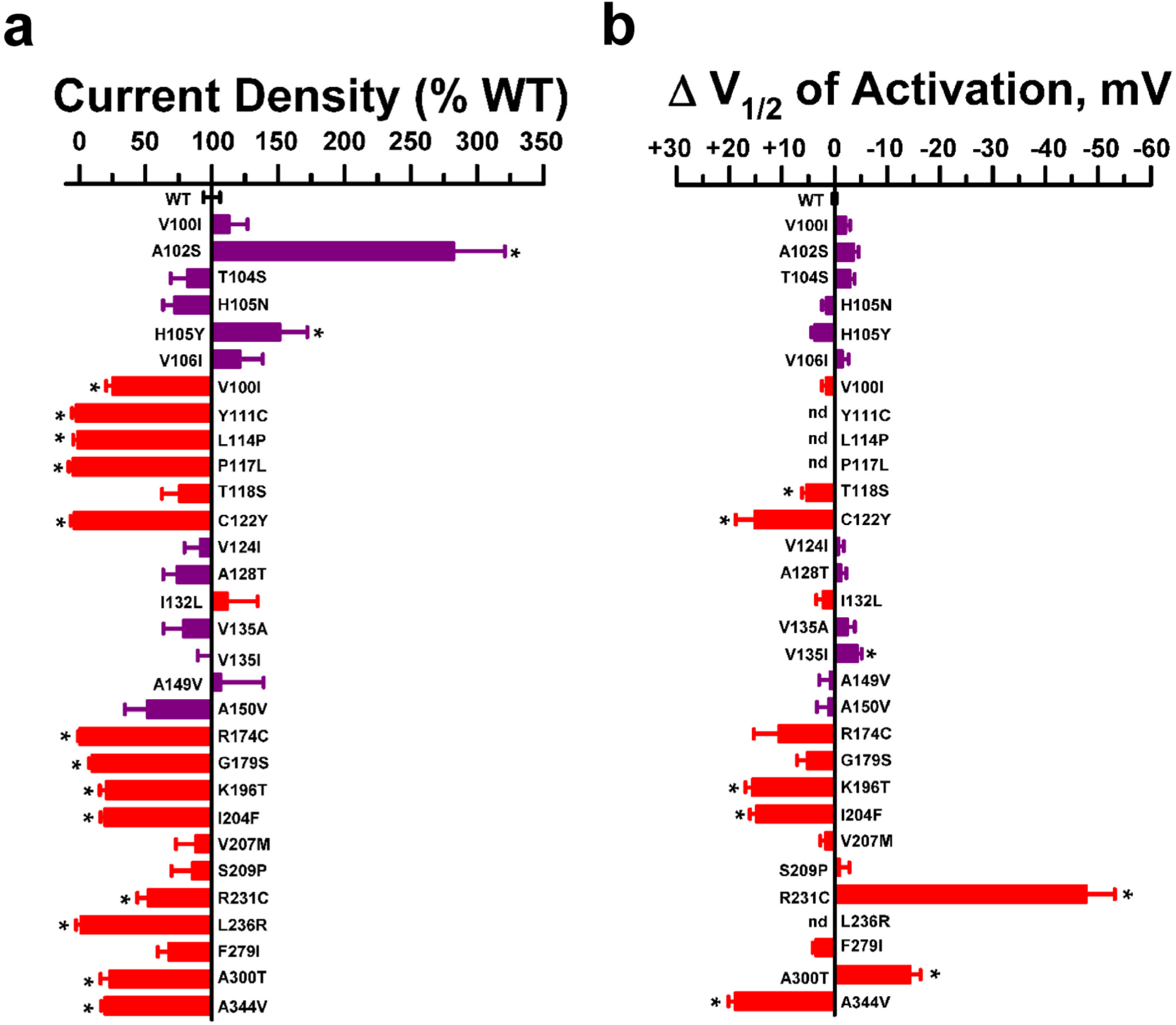
Functional properties of training set KCNQ1 variants recorded by automated patch clamp. **a**) Average whole-cell currents recorded at +60 mV from cells expressing training set KCNQ1 variants plotted as percent of WT (n = 23-137). **b**) Difference in activation V_½_ relative to WT determined for variants with current amplitude >10% of WT (n = 4–79). KCNQ1 variants are ordered by codon number with red bars indicating disease-associated variants and magenta bars representing non-disease variants. Bar height indicates mean value and error bars are SEM (* indicates p –0.01).

**Fig. 4.**
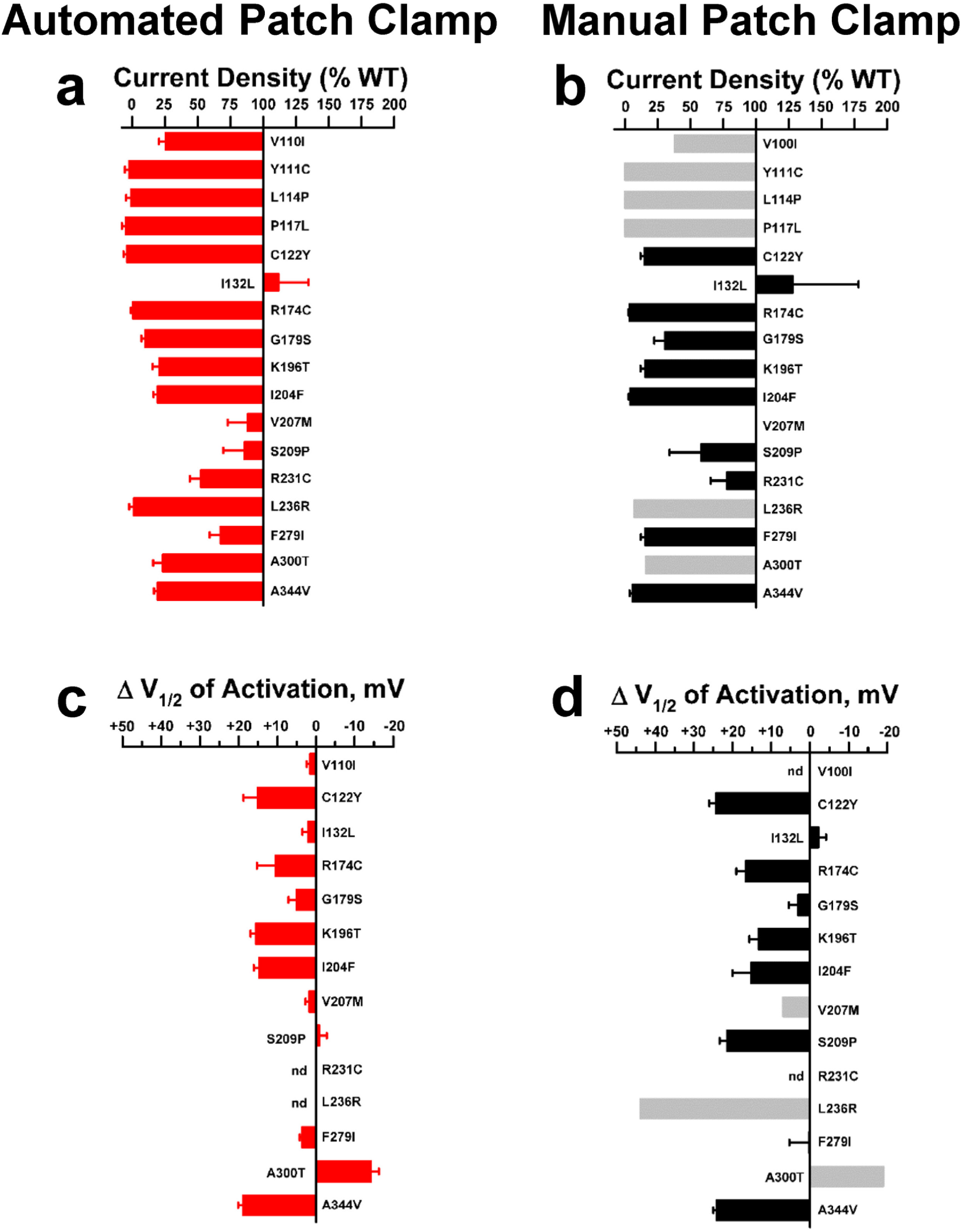
Validation of automated patch clamp recording for KCNQ1 variants. **a,b**) Average whole-cell current density for training set disease-associated KCNQ1 variants determined by automated (*left*) or manual (*right*) patch clamp recording measured at +60 mV and expressed as percent of WT. For manual patch clamp, grey shaded bars are values derived from published reports (mean values only, see below for citations) and black shaded bars are data from this study (mean, SEM). Quantified functional parameters are provided in Supplemental Tables S2a and S2b. **c,d**) Differences in activation V_½_ relative to WT for training set disease-associated KCNQ1 variants (nd = not determined). Functional data from the literature are for V100I,^34^ Y111C,^35^ L114P,^35^ P117L,^35^ V207M,^36^ L236R,^37^ and A300T.^38^

We also quantified the time course of deactivation by fitting current traces with exponential functions (fit parameters given in Supplemental Table S2a). Variant R231C is notable for a markedly slower time course of deactivation compared with WT channels (Fig. 5) and this resembles published data obtained using manual patch clamp recording.^29^ Impaired deactivation was also observed for variants I132L and S209P, while faster deactivation was exhibited by K196T. Similar results were observed by manual patch clamp recording in our laboratory (Supplemental Table S2b). None of the other functional training variants exhibited abnormal gating kinetics and this is consistent with literature reports.

**Fig. 5.**
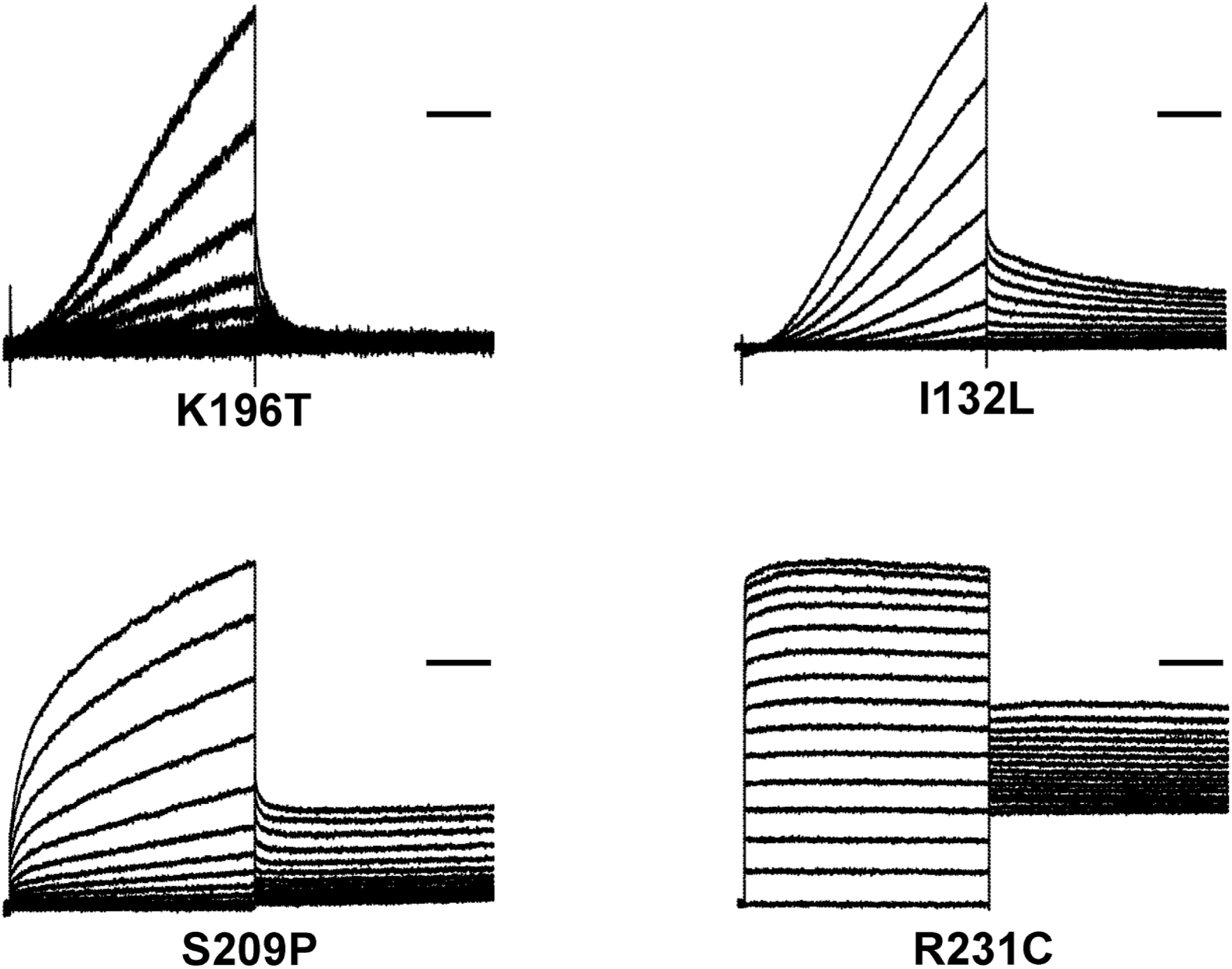
Altered gating kinetics for KCNQ1 variants detected by automated patch clamp. Average whole-cell currents recorded from cells expressing select KCNQ1 variants to illustrate the capability of automated patch clamp to detect altered gating kinetics. Traces were normalized to peak current measured at +60 mV to enable comparison of gating behaviors among variants with divergent current density. Horizontal bars represent 500 ms. Time constants of deactivation are presented in Supplemental Table S2a.

**Fig. 6.**
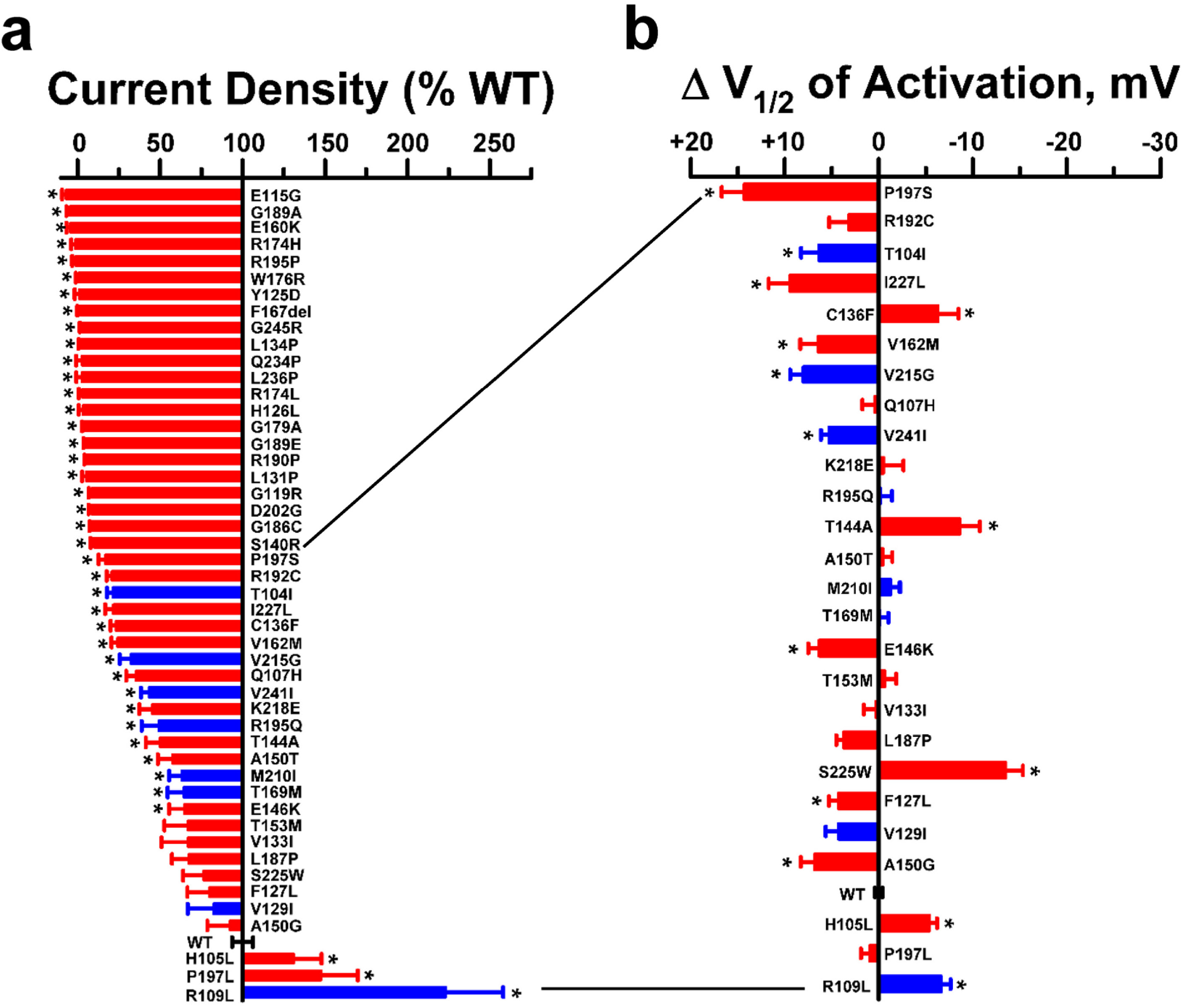
Functional properties of KCNQ1 variants of unknown significance. **a**) Plot of average whole-cell current density (measured at +60 mV) recorded from cells expressing test KCNQ1 variants and displayed as percent of WT (n = 23-94). KCNQ1 variants are ordered from lowest to greatest current density with red bars indicating disease-associated variants and blue bars representing ExAC variants. **b**) Difference in activation V_½_ relative to WT determined for variants with current amplitude >10% of WT (n = 6-55). Variants are listed in the same order as in panel **a**. Bar height indicates mean value and error bars are SEM (* indicates p ≤ 0.015). Quantified functional parameters are provided in Supplemental Tables S2c.

### Functional properties of KCNQ1 variants of unknown significance

We used this optimized approach to determine the functional properties of 48 additional KCNQ1 variants that have not been studied previously (Supplemental Table S3). Sixteen disease-associated variants were from the Human Gene Mutation Database (HGMD)^22^ and an additional 24 variants of unknown significance were provided in de-identified formats by two commercial genetic testing laboratories (GeneDx, Inc., Gaithersburg, MD; Transgenomic Inc., New Haven, CT). The associated disease was LQTS for the majority of the variants. Eight additional rare variants were from the Exome Aggregation Database (ExAC).^30^

Recordings were made from 12 separate transfection experiments (average transfection efficiencies were KCNQ1: 84 ± 6.7%, KCNE1: 79.4 ± 7.6% and co-transfection: 75 ± 7.9%; with viability of 94.9 ± 4.7%) and data from 2,566 cells were used for final analysis (12 384-well plates, average 46 cells per variant; n = 343 WT cells). Figure 6a illustrates a waterfall plot of average current density for each variant expressed as percent of WT. Representative whole-cell current traces for each variant are presented in Supplemental Fig. S4. The majority of the disease-associated variants and two rare variants from ExAC exhibited severe loss-of-function (defined as ≤ 25% of WT current density). The remaining variants exhibited more modest differences in current density compared with WT channels with the exception of R109L, a rare ExAC variant that exhibited a profound gain-of-function (223% of WT). Two other ExAC variants (T104I, V215G) exhibited significantly lower current density than WT channels (22% and 35% of WT current, respectively; p < 0.015 for both) demonstrating that rare variants present in population databases can have functional effects.

Some of the variants in the test set with preserved current density exhibited effects on the voltage-dependence of activation or had altered gating kinetics including slower (e.g., T144A and S225W) or faster (e.g., C136F and V244I) deactivation (Fig. 6b; Supplemental Fig. S5; Supplemental Table S2c). Notable is the behavior of P197L, a rare variant originally described in a study of cost effectiveness of genetic testing for heritable cardiac disorders including LQTS,^31^ but also discovered incidentally in a breast cancer cohort undergoing exome sequencing.^32^ This variant exhibits a moderate gain-of-function (145% of WT current) that is inconsistent with a mutation causing LQTS. This finding highlights the value of experimental evidence for accurate variant classification.

### Correlation of variant allele frequency with channel function

Among the *KCNQ1* variants studied, 20 were present in ExAC with minor allele frequencies less than 0.0003 (rare variants) whereas 32 of the test variants were absent from this population exome database (ultra-rare variants). We examined the relationship of minor allele frequency with channel function for rare and ultra-rare variants. As illustrated in Fig. 7, there was no correlation between the rarity of KCNQ1 variants and the measured current density. This observation suggests that allele frequency is not a reliable predictor of variant channel function.

**Fig. 7.**
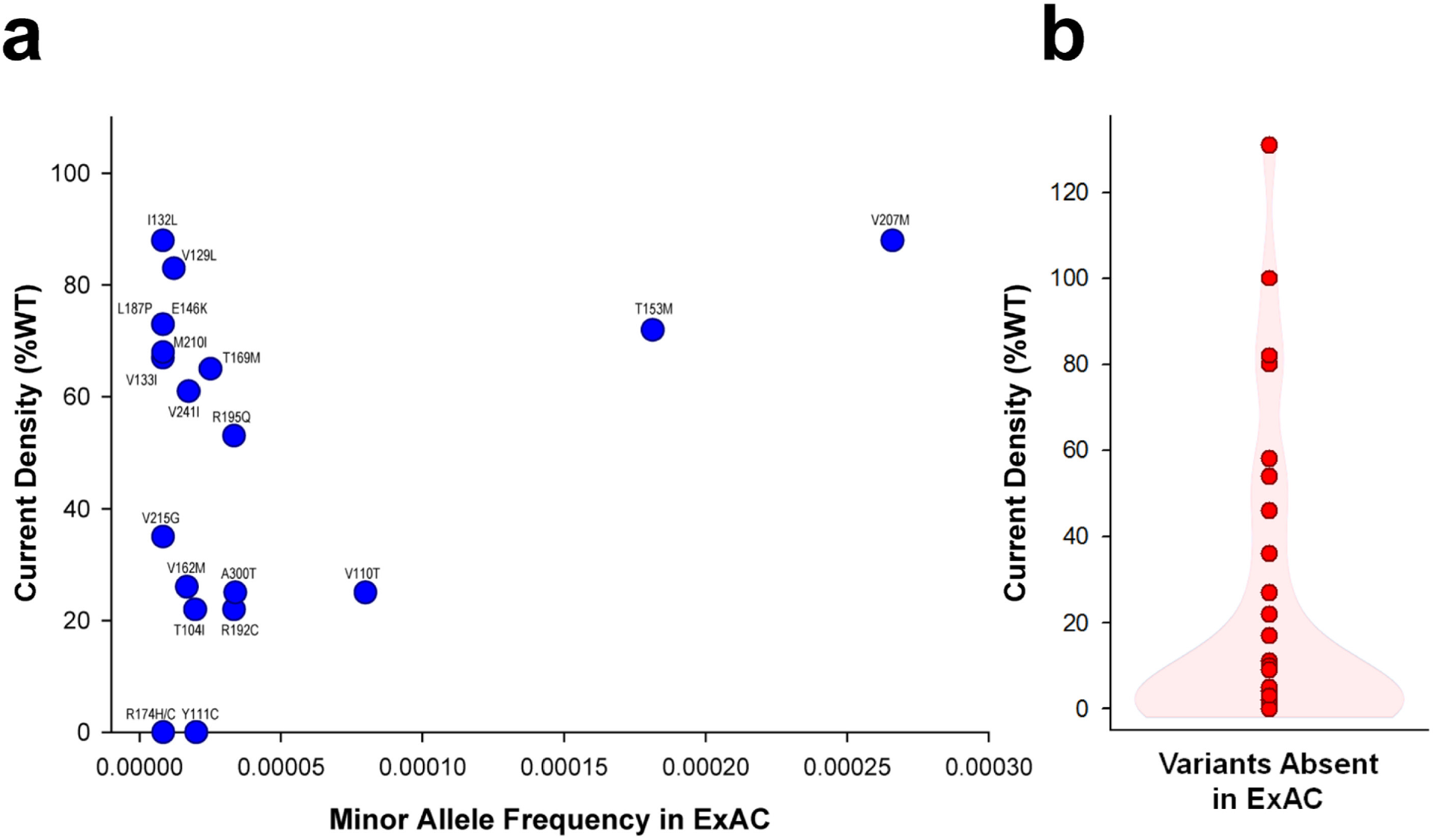
Correlation of allele frequency and channel function for rare KCNQ1 variants. **a**) Current density expressed as percent of WT for a group of KCNQ1 variants with minor allele frequencies <0.0003 in ExAC. Individual variant names are placed beside each data symbol. **b**) Current density for 32 KCNQ1 variants that are not found in ExAC. A violin plot of the dataset is superimposed on the individual data points.

### Impact of functional evaluation on variant classification

We assessed the potential impact of our functional data on the classification of *KCNQ1* variants. Among the 78 variants investigated in this study, 39 were annotated in the ClinVar database with assertions regarding the likelihood of pathogenicity ranging from no assertion to pathogenic. Using a conservative approach, we reclassified those variants according to their functional phenotype. Variants with severe loss-of-function (≤ 25% of WT) were reclassified as ‘likely pathogenic’ when the original ClinVar classification was uninformative (e.g., VUS, conflicting interpretations, no assertion provided), or as ‘pathogenic’ if originally classified as ‘likely pathogenic’. Variants with normal or near normal functional properties were reclassified as ‘likely benign’ when there was no assertion or conflicting interpretations. Finally, variants with milder degrees of dysfunction were classified as VUS to reflect uncertainty regarding pathogenicity.

Using this conservative reclassification strategy, 21 of 29 (72%) variants originally classified in uninformative categories could be reclassified as either ‘likely pathogenic’ or ‘likely benign’ (Fig. 8). All 6 variants classified as ‘likely pathogenic’ in ClinVar could be reclassified as ‘pathogenic’ because of severe loss-of-function, whereas all of the variants originally categorized as ‘pathogenic’ remained in this category. Our analysis also included 6 variants (H105L, I132L, E146K, A150G, L187P, V207M) associated with LQTS that could be reclassified as ‘benign’ or ‘likely benign’ based on near normal channel function. Overall, our functional evaluation could impact the classification of ~70% of KCNQ1 variants annotated in ClinVar.

**Fig. 8.**
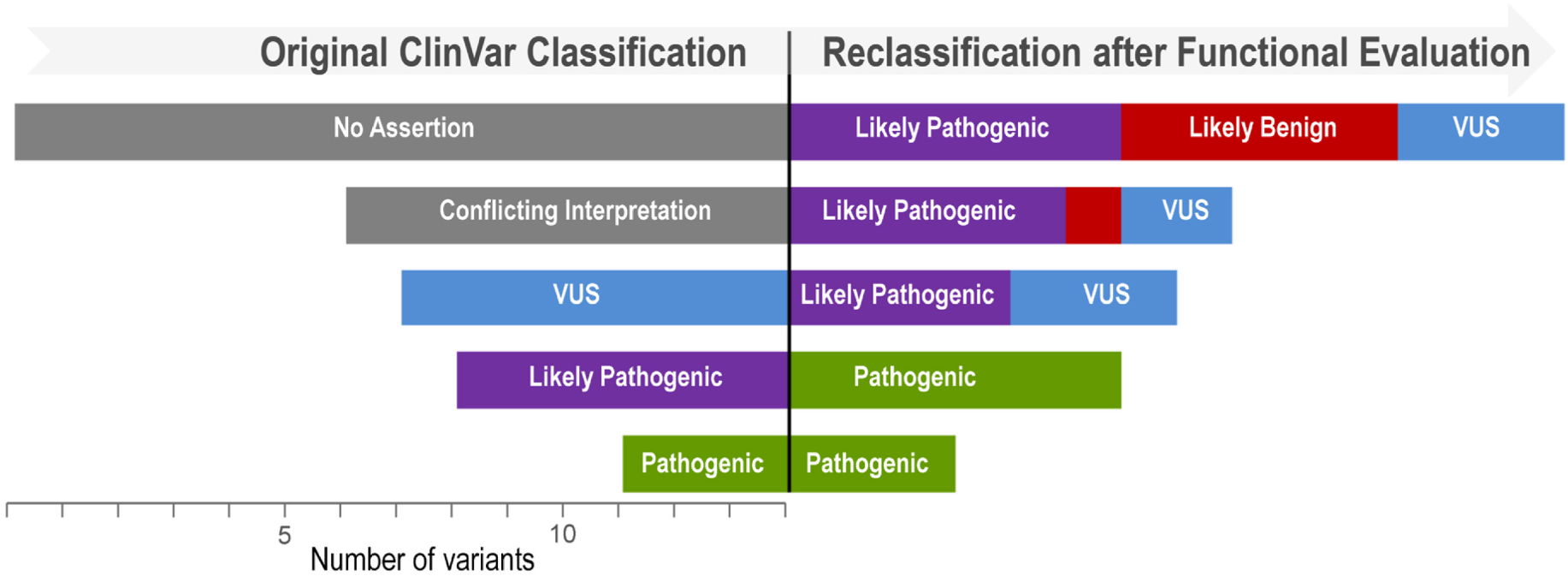
Reclassification of KCNQ1 variants. The original classification in ClinVar is represented on the left half of the display and the reclassification based on the functional data is illustrated by the right half. The horizontal bars represent the variants classified in different categories and the length of the bars is proportional to the number of variants. Original classifications: No assertion, n = 14; Conflicting Interpretation, n = 8; VUS, n = 7; Likely Pathogenic, n = 6; Pathogenic, n = 3.

## DISCUSSION

Mutations in genes encoding ion channels are associated with a variety of monogenic disorders with diverse clinical manifestations. The emergence of widespread clinical genetic testing and the use of next-generation sequencing in genetic research has resulted in explosive growth of known ion channel variants associated with disease traits and in populations. As with other genetic disorders for which genetic testing is routine, discriminating pathogenic from benign variants and establishing genotype-phenotype relationships has become increasingly more challenging as the number of variants grows.

A new and widely embraced scheme for variant classification proposed by the American College of Medical Genetics and Genomics (ACMG) is based on sets of weighted criteria that can be scored to assess the likelihood of pathogenicity.^3^ Among the criteria weighted as strongly supportive of classifying variants as pathogenic is the availability of data from *in vitro* or *in vivo* functional studies demonstrating a damaging effect on the gene or gene product (PS3 criterion).^3^ By contrast, *in silico* predictions of variant effects are considered supporting evidence, the least strong criterion. Our approach to high throughput functional evaluation of *KCNQ1* variants offers a rich new data source that satisfies the PS3 criteria and can aid variant classification. Importantly, although our findings for many of the LQTS-associated variants are consistent with severe loss-of-function and consequent pathogenesis of the disease, our data are best viewed as informing about the functional properties of variants and not a judgement about pathogenicity. Final assertions about variant pathogenicity need also to consider phenotypic and genetic data. However, our conceptual experiment assessing the potential impact of functional data on variant classification (Fig. 8) suggests that a large proportion of variants with uninformative or no assertions of pathogenicity could be moved into more definitive categories.

A major technological advance enabling modern studies of ion channels has been patch clamp electrophysiology, which underlies decades of important discoveries in several disciplines. For channelopathies, *in vitro* functional assessments of ion channel function using patch clamp recording has been the cornerstone of research determining the functional impact of mutations and establishing genotype-phenotype relationships. Although the patch clamp technique brought about a revolution in ion channel biology, the method in its typical embodiment has limited throughput and is extremely time and labor intensive. In an abstract sense, ion channel electrophysiology has advanced in a manner similar to that of DNA sequencing technology, although its democratization has lagged far behind sequencing. Originally, DNA sequencing was a difficult, low-throughput method performed only in elite genetics and molecular biology laboratories having specialized expertise and equipment.^33^ As methods became more standardized, more laboratories adopted DNA sequencing to perform small scale projects. Development of automated sequencing instruments enabled more widespread use and larger scale projects such as genome sequencing. Similarly, in the early days of patch clamp electrophysiology, methods were difficult and non-standardized with only a few laboratories having the necessary expertise and requisite equipment. Over time, standardization of hardware and software allowed more investigators to perform patch clamp recording for an expanding array of research applications. Newly developed automated patch clamp instruments have been implemented primarily for drug discovery, but these technologies are poised to enable wider applications including the large scale functional annotation of genetic variants in ion channel genes, as we demonstrated in this study.

In this study, we utilized automated patch clamp recording to enable a large scale functional analysis of *KCNQ1* variants. To enable effective use of this technology, we first optimized a highly efficient method for transient transfection of heterologous cells using a high capacity electroporation platform. The unique combination of these two technologies provided a strategy with a much greater throughput than standard methods. We validated this approach by comparing results with manual patch clamp recording including published data from other laboratories. We elucidated the functional consequences of 48 variants of unknown significance in *KCNQ1* and this represents ~50% increase in the number of variants with known function. Importantly, the bulk of the experimental work involved with studying the 48 *KCNQ1* variants was completed in approximately 12 weeks and additional parallelization of the workflow can further accelerate this timeline. In addition to demonstrating the functional effects of several *KCNQ1* variants, which alone will likely contribute to classifying many variants, we also revealed that the rarity of allele frequency in reference populations is not a reliable indicator of channel dysfunction. Efficient, high-throughput experimental functional studies of ion channel variants can be harnessed to decrypt many more variants of unknown significance.

In summary, we demonstrated the feasibility and robustness of a novel union of technologies (high efficiency cell electroporation, automated planar patch clamp recording) in a scalable platform for rapidly assessing the functional consequences of genetic variants in a human ion channel gene, *KCNQ1*. The workflow can be adapted for assaying variants in other ion channels and should also be suitable for pharmacological profiling of individual variants. Elucidating the function of human ion channel variants using this high throughput experimental approach can promote data-driven variant classification and create new opportunities for precision medicine.

## ACKNOWLEDGEMENTS

The authors thank Dr. Tom Callis for providing KCNQ1 variant data from Transgenomic, Inc.

## AUTHOR CONTRIBUTIONS

C.R.S., C.G.V and A.L.G. conceived and designed the study; R.R.D. and K.L.F. performed mutagenesis, cell culture and electroporations; C.G.V. and F.P. performed electrophysiological recordings and data analysis; J-M.D. wrote custom data analysis software; D.M. provided information on unpublished variants; J.M. provided the literature review of variant function; C.R.S., J.M. and A.L.G. acquired funding for the project. C.G.V. and A.L.G. wrote the manuscript with input from all authors.

